# Reconstitution of muscle cell microtubule organization in vitro

**DOI:** 10.1101/2022.01.26.477920

**Authors:** Ambika V. Nadkarni, Rebecca Heald

## Abstract

Skeletal muscle differentiation occurs as muscle precursor cells (myoblasts) elongate and fuse to form multinucleated syncytial myotubes in which the highly-organized actomyosin sarcomeres of muscle fibers assemble. Although less well characterized, the microtubule cytoskeleton also undergoes dramatic rearrangement during myogenesis. The centrosome-nucleated microtubule array found in myoblasts is lost as the nuclear membrane acquires microtubule nucleating activity and microtubules emerge from multiple sites in the cell, eventually rearranging into a grid-like pattern in myotubes. In order to characterize perinuclear microtubule organization using a biochemically tractable system, we isolated nuclei from mouse C2C12 skeletal muscle cells during the course of differentiation and incubated them in cytoplasmic extracts prepared from eggs of the frog *Xenopus laevis*. Whereas centrosomes associated with myoblast nuclei gave rise to radial microtubule arrays in extracts, myotube nuclei produced a sun-like pattern with microtubules transiently nucleating from the entire nuclear envelope. Perinuclear microtubule growth was suppressed by inhibition of Aurora A kinase or by degradation of RNA, treatments that also inhibited microtubule growth from sperm centrosomes. Myotube nuclei displayed microtubule motor-based movements leading to their separation, as occurs in myotubes. This in vitro assay therefore recapitulates key features of microtubule organization and nuclear movement observed during muscle cell differentiation.

## INTRODUCTION

The cytoskeleton of muscle cells is highly organized and specialized with actin, myosin, and many associated proteins forming contractile sarcomeres (Henderson et al., 2017). Much less is known about microtubules, which arrange into grid-like arrays parallel to the long axis of sarcomeres in myofibers, the giant multinucleated cells of skeletal muscle, where they are thought to provide viscoelastic resistance, maintain nuclear integrity in the presence of vigorous contractile forces, and contribute to proper spacing of nuclei at the cell periphery (Azevedo & Baylies, 2020; Becker et al., 2020; Caporizzo et al., 2018; Heffler et al., 2020; Oddoux et al., 2013; Robison et al., 2016; Warren, 1974). Defects in microtubule associated proteins are related with nuclear mis-positioning that can impede muscle function, but the molecular mechanisms leading to the organization and function of the microtubule cytoskeleton in muscle cells remain poorly understood (Azevedo & Baylies, 2020).

The microtubule cytoskeleton is dramatically reorganized during muscle cell differentiation (Abmayr & Pavlath, 2012; Becker et al., 2020). Microtubule depolymerization and re-growth assays have demonstrated that microtubules in differentiated muscle cells do not nucleate from centrosomes, but polymerize from non-centrosomal microtubule organizing centers (ncMTOCs) found in the cytoplasm and on the nuclear envelope (Gimpel et al., 2017; Kronebusch & Singer, 1987; Musa et al., 2003; Oddoux et al., 2013; Tassin et al., 1985). A key step in microtubule reorganization in these cells involves the expression of specific isoforms of the nesprin family of outer nuclear-membrane spanning, nucleoskeletal proteins (Apel et al., 2000; Duong et al., 2014; Randles et al., 2010; Zhang et al., 2001). In cultured muscle cells, Nesprin 1α recruits microtubule organizing proteins to the nuclear envelope, such as AKAP450, PCM-1 and pericentrin, three proteins that are also important for centrosomal microtubule nucleation (Balczon et al., 1994; Dammermann & Merdes, 2002; Dictenberg et al., 1998; Doxsey et al., 1994; Espigat-Georger et al., 2016; Gimpel et al., 2017; Kubo et al., 1999; Schmidt et al., 1999; Srsen et al., 2009; Takahashi et al., 1999; Witczak et al., 1999). Microtubule-based motors dynein and kinesin also localize to the nuclear envelope where they mediate nuclear movements (Azevedo & Baylies, 2020; Cadot et al., 2012; Wilson & Holzbaur, 2012, 2015).

A challenge in the field is to understand the molecular basis of microtubule rearrangements during myogenesis, including how centrosomal microtubule nucleation is attenuated, and how ncMTOCs arise. Indeed, non-centrosomal microtubule organization observed in myotubes is a common feature of many differentiated cell types that reorganize their microtubule cytoskeletons to serve particular functions, such as establishing the polarity of epithelial cells (Sanchez & Feldman, 2017). There is also a need to elucidate precisely how microtubule populations emerging from various nucleation centers in the cell contribute to nuclear positioning in animals, thereby revealing potential therapeutic targets for muscular dystrophies, as well as novel, yet undiscovered roles for microtubules and associated factors at the nuclear envelope.

To begin addressing these questions, we developed an assay based on *Xenopus laevis* egg extracts that recapitulate interphase cytoplasmic organization around nuclei in vitro, and the mouse C2C12 culture system in which precursor myoblast cells differentiate into syncytial myotubes that localize ncMTOCs to the nuclear membrane and form ordered actomyosin arrays (Bugnard et al., 2005; Cheng & Ferrell, 2019; Lu et al., 2001; Musa et al., 2003). By isolating nuclei at different stages of muscle cell differentiation and incubating them in egg extract, we could reproduce features of myoblast and myotube microtubule organization (Fant et al., 2009; Lu et al., 2001; Musa et al., 2003; Srsen et al., 2009). Whereas myoblast nuclei produced centrosome-nucleated arrays, myotube nuclei generated distinct sun-like patterns of perinuclear microtubules, indicating robust microtubule nucleation activity at the nuclear membrane. We show that microtubule polymerization from the nuclear envelope requires Aurora A kinase activity and is inhibited by RNase treatment, and that muscle cell nuclei move apart in a microtubule- and motor-dependent manner. These results show that important aspects of muscle-specific microtubule organization can be reconstituted in vitro, providing a novel assay for analyzing mechanisms of ncMTOC function in differentiating cells.

## RESULTS AND DISCUSSION

### Purified muscle nuclei retain the ability to nucleate and organize microtubules

Nuclei isolated from mouse skeletal muscle C2C12 cells differentiated in culture for 4-5 days were added to *Xenopus laevis* egg extracts in interphase of the cell cycle and supplemented with rhodamine-labeled tubulin. Within 10 minutes, microtubules were observed emanating from the surface of nuclei in a sun-like array (Figure 1A), similar to the pattern seen in mono-nucleated, differentiating muscle cells in culture (Srsen et al., 2009). Perinuclear microtubule fluorescence intensity decreased gradually with increasing distance from each nucleus (Figure 1B) and a very weak correlation between nuclear area and total microtubule intensity was observed (Figure 1C) (R^2^= 0.045, p=0.02). To determine whether the in vitro system recapitulated differences between undifferentiated and differentiated C2C12 cells, the assay was performed with nuclei isolated from successive stages of differentiation. Whereas undifferentiated myoblasts (Day 0) nucleated microtubules from a focal centrosome adjacent to the nucleus, myotube nuclei (Day 5) organized microtubules all around the nuclear periphery (Figure 1D). Similar perinuclear microtubule organization was observed with nuclei isolated from cultures day 2-6 post differentiation, but after 7 days individual nuclei became difficult to isolate and observe, as syncytial cells grew larger and nuclei clumped together during preparation (unpublished data).

**Figure 1.**
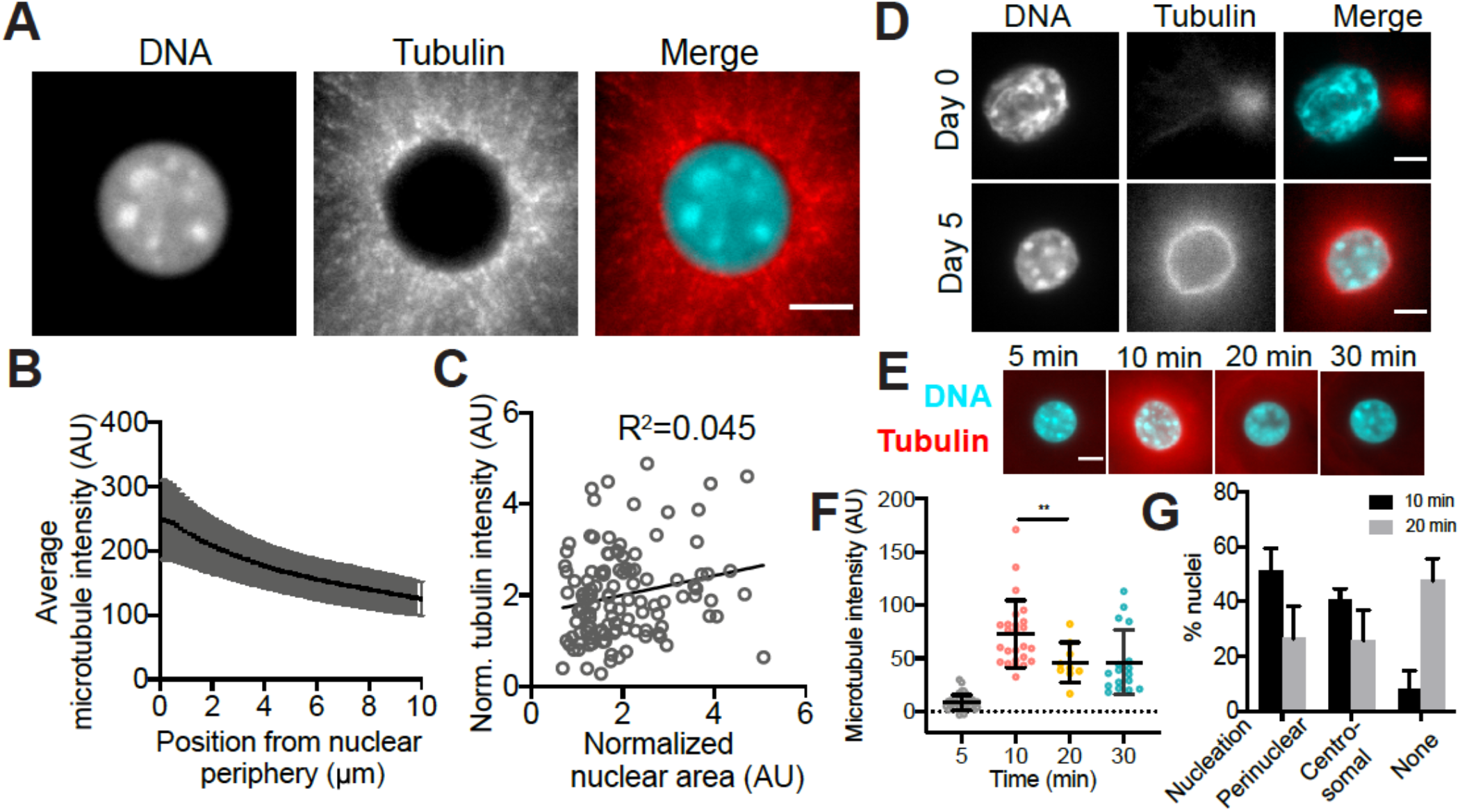
Differentiated C2C12 nuclei organize microtubules in *Xenopus* egg extract. (A) Fluorescence images of microtubules (red) radiating from C2C12 nuclei (cyan) isolated from differentiated myotubes incubated in *Xenopus* egg extract. (B) Microtubule fluorescence profile of a 10 µm zone from the edge of each nucleus shows decreasing intensity from the nuclear periphery. (C) Plot of normalized nuclear area and total microtubule intensity shows a very slight positive correlation (R^2^= 0.045, p=0.02, n = 3 extracts). (D) Comparison of microtubule polymerization from undifferentiated (Day 0) and differentiated (Day 5) nuclei illustrate the change in microtubule nucleation patterns. (E) Representative images of microtubules around myotube nuclei fixed in squashes at 5, 10, 20 and 30 minutes after addition to *Xenopus* egg extract. (F) Corresponding quantification showing that fluorescence of microtubules proximal to the nuclear surface peaks at 10 minutes (representative extract out of n=3 is depicted, p= 0.0065). (G) Quantification of nucleation activity at 10 and 20 minutes shows that the percentage of nuclei with centrosomal or perinuclear nucleation decreases over time as the percentage nuclei that exhibit no microtubule nucleation increases by 50%. Scale bars are 5 µm.

A time-course of microtubule polymerization from C2C12 nuclei in egg extracts revealed that nucleation in a 5 µm zone proximal to the nuclear membrane was most intense at 10 minutes of incubation and decreased significantly by 20 minutes (from 72.8±31.7 AU to 46.2±18.8 AU) (Figure 1E and F) with a lower proportion of nuclei showing either perinuclear or centrosomal microtubule nucleation at later time points (Figure 1G). We therefore used 10 minutes as the endpoint in subsequent assays.

This in vitro system therefore demonstrates that nuclei from muscle cells retain microtubule organization capacity corresponding to their differentiation state as observed in cultured cells (Fant et al., 2009; Kronebusch & Singer, 1987; Srsen et al., 2009; Tassin et al., 1985). However, our assay did not detect a relationship between ncMTOC area and total microtubule intensity, perhaps because microtubule polymerization was variable and/or our assay is not sensitive enough. The transient nature of nuclear envelope-associated microtubule nucleation is interesting and consistent with the pattern of microtubule growth in differentiating muscle, as perinuclear microtubules are thought to dissociate and rearrange into bundles parallel to the long axis of the cell (Figure 2A) (Lu et al., 2001; Oddoux et al., 2013; Tassin et al., 1985; Warren, 1974). Analogous parallel microtubule organization was not observed in egg extracts at later time points, which is not surprising considering differences in composition and organization of the cytoplasm compared to muscle cells. Future experiments will investigate mechanisms underlying the burst of nuclear envelope-associated ncMTOC activity observed both in vivo and in vitro.

**Figure 2.**
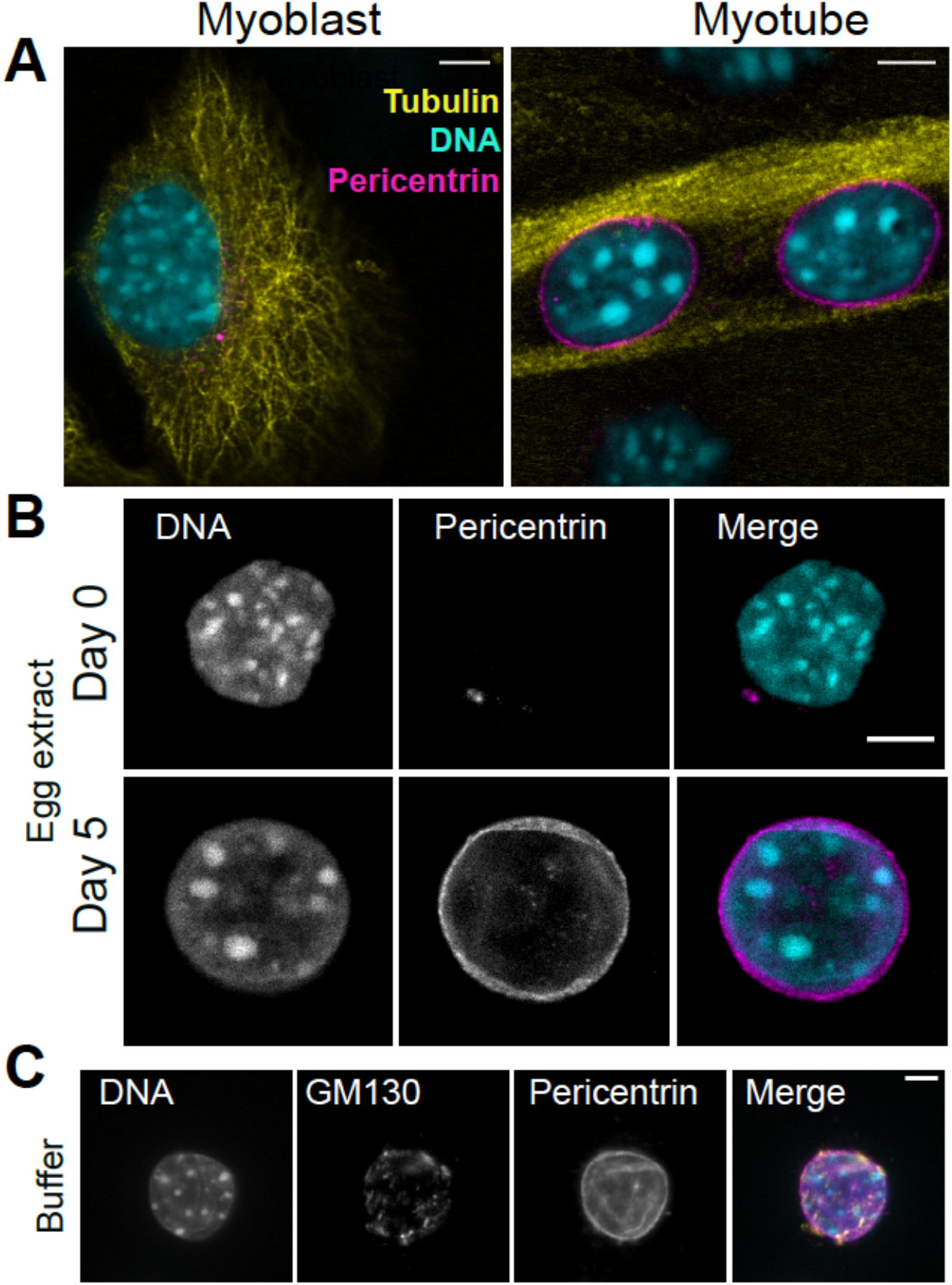
Pericentrin redistributes to the nuclear membrane of myotube nuclei and remains associated with isolated muscle cell nuclei. (A) Immunofluorescence images showing that pericentrin (magenta) localizes to the centrosome in an undifferentiated C2C12 myoblast and to the nuclear envelope in a differentiated myotube, where microtubules (yellow) have reorganized parallel to the long axis of the syncytial cell. (B) Images of isolated myoblast and myotube nuclei incubated in *Xenopus* egg extract exhibit similar pericentrin localization to the centrosome and nuclear periphery, respectively. (C) Golgi elements labeled with antibodies to GM130 (yellow in merged image) associate with nuclei purified from C2C12 myotubes and incubated in buffer, partially overlapping with pericentrin at the nuclear periphery. Scale bars are 5 µm.

### Microtubule-organizing factors stably associate with muscle cell nuclei

To examine the source of the microtubule nucleation patterns observed in the in vitro system, reactions were fixed and spun down onto coverslips and examined by immunofluorescence for the centrosome scaffolding protein pericentrin (Dictenberg et al., 1998; Doxsey et al., 1994). As observed in cultured myoblasts and myotubes (Figure 2A), pericentrin colocalized with sites of microtubule nucleation at centrosomes attached to myoblast nuclei, or at the myotube nuclear periphery (Figure 2B)(Musa et al., 2003). Isolated myotube nuclei added to buffer instead of egg extract exhibited similar pericentrin localization (Figure 2C), indicating a stable association with the nuclear envelope during preparation as previously described (Srsen et al., 2009).

The Golgi complex, which is also recognized as an ncMTOC, surrounds myotube nuclei and was recently reported to be a major source of microtubule nucleation in differentiating skeletal muscle cells based on the localization of the centrosome- and Golgi-associated protein CDK5RAP2 at sites of perinuclear microtubule growth and the inhibitory effects of brefeldin A treatment (Chabin-Brion et al., 2001; Ide et al., 2021; Kronebusch & Singer, 1987; Oddoux et al., 2013; Ralston, 1993). We therefore stained isolated myotube nuclei using antibodies to GM130, a Golgi matrix component, and observed puncta that partially colocalized with pericentrin (Figure 2C). To determine if Golgi elements were necessary for perinuclear ncMTOC activity, we attempted to dissociate them by treating isolated nuclei with higher concentrations of detergent, but this also disrupted the nuclear envelope and prevented further analysis. Neither treatment of C2C12 cells with brefeldin A and subsequent purification of nuclei, nor direct treatment of egg extract with brefeldin A, yielded reproducible effects on perinuclear microtubules compared to controls (unpublished data).

Altogether, these observations indicate that pericentrin and other factors mediating microtubule polymerization remain associated with mouse muscle cell nuclei during preparation. To what extent Golgi elements play a role in perinuclear microtubule growth in egg extracts remains to be determined.

### Aurora A and RNA are required for perinuclear microtubule growth

We next used the in vitro system to identify molecules important for perinuclear microtubule nucleation and to test whether conserved pathways control both centrosomal and perinuclear microtubule polymerization in egg extracts. We first evaluated the Aurora A kinase, which functions in centrosome maturation, recruitment of pericentriolar material, and microtubule nucleation (Magnaghi-Jaulin et al., 2019). Beads coated with Aurora A act as ncMTOCs, however a role for this kinase in perinuclear microtubule nucleation has not been described (Ishihara et al., 2014; Nguyen et al., 2014; Tsai & Zheng, 2005). Treatment of *Xenopus* egg extracts with the Aurora A inhibitor MLN8237 greatly diminished microtubule nucleation around C2C12 nuclei (Figure 3A and B) and nearly abolished microtubule nucleation at sperm centrosomes (Figure 3C and D), indicating that similar mechanisms underlie microtubule nucleation in both cases in egg extract. To test whether Aurora A inhibition diminished perinuclear microtubule nucleation by displacing pericentrin, we measured its intensity around myotube nuclei and found it was unchanged in control versus drug-treated nuclei (Supplementary Figure S1).

**Figure 3.**
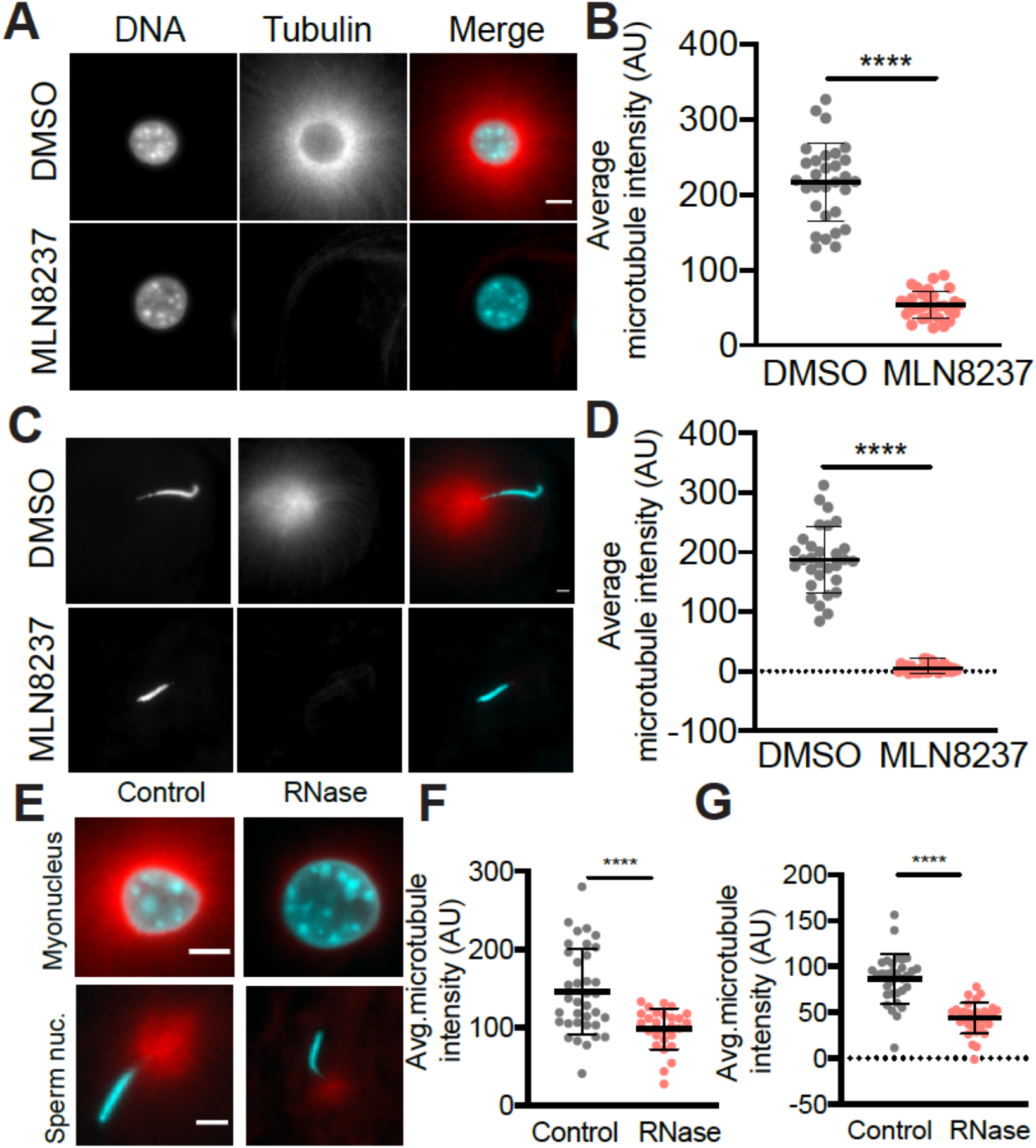
Perturbation of Aurora-A kinase activity and degradation of RNA inhibit microtubule growth around both myotube nuclei and sperm centrosomes. (A) Representative fluorescence images of myotube nuclei (cyan) in *Xenopus* egg extract treated with 0.5% DMSO or 1 µM Aurora A inhibitor MLN8237 show that microtubules (red) are greatly reduced in the presence of the drug. Scale bar is 5 µm. (B) Plot of quantified average microtubule fluorescence of a 5 µm zone around the nuclear periphery shows that microtubule assembly is significantly decreased upon kinase inhibition (control = 216.4, S.D.=51.77, drug-treated = 53.71, S.D.=18.37, p < 0.0001 (t-test with Welch’s correction)). Graph shows a representative extract from 3 independent experiments, n >= 30 nuclei for each. (C) Representative images of sperm nuclei in DMSO- and MLN8237-treated extracts show that the robust microtubule aster normally emanating from the sperm centrosome is lost upon kinase inhibition. Scale bar = 2 µm (D) Plot of quantified average microtubule fluorescence of a fixed area of radius 20 µm shows that microtubule assembly from centrosomes is strongly impaired (control = 187.2, S.D.=54.81, drug-treated = 5.83, S.D.=7.49, p < 0.0001 (t-test with Welch’s correction)) (E) Representative images of myotube nuclei or sperm centrosomal asters in extract treated with buffer or 100 µg/ml RNase-A that shows reduced microtubule nucleation. Scale bar = 5 µm. (F) Quantification of average microtubule intensity shows that microtubule nucleation is impaired around myotube nuclei (control = 146.3, S.D.= 54.55, RNase-treated = 98.32, S.D.=26.30, p < 0.0001 (t-test with Welch’s correction)) as well as (G) sperm centrosomes when extract is treated with RNase-A (control = 86.45, S.D.=27.16, RNase-treated = 43.66, S.D.=16.86, p < 0.0001 (t-test with Welch’s correction)).

Previous work has shown that RNA localizes to microtubules and that RNA-containing protein complexes (RNPs) function to stabilize spindle microtubules in metaphase *Xenopus* egg extracts (Blower et al., 2005). We found that treating interphase egg extracts with RNase A similarly impaired microtubule polymerization, reducing microtubule growth at both myotube nuclei (∼1.5 fold) and sperm centrosomes (∼2-5 fold) (Figure 3E-G). Co-translational targeting of pericentrin mRNA has been shown to contribute to centrosome maturation (Sepulveda et al., 2018). However, because perinuclear microtubule nucleation occurs even in the presence of translation inhibitors (unpublished data), we favor a translation-independent model in which RNA contributes directly to microtubule stabilization by promoting formation of RNPs such as the Rae1 complex, which was shown to be regulated by importin β and RanGTP in metaphase egg extracts (Blower et al., 2005). To address a potential role for this type of regulatory mechanism, we added recombinant Ran mutants Q69L or T24N to activate or inhibit the pathway, respectively (Cavazza & Vernos, 2015), but did not observe significant effects on perinuclear microtubule growth (unpublished data).

Overall, these findings support a model in which common factors control microtubule nucleation at centrosomes and at the periphery of myotube nuclei in the interphase egg cytoplasm, including Aurora A and its substrates as well as microtubule-stabilizing RNPs. In contrast, a major pathway controlling ncMTOC activity in metaphase that depends on RanGTP, does not appear to operate. Perturbations combining RNAi of specific factors in C2C12 myotubes prior to nuclear isolation and depletion of candidate microtubule organizing proteins from egg extracts will be essential to identify and characterize the key drivers of perinuclear microtubule nucleation and growth in this assay.

### Movement of myotube nuclei in vitro is driven by microtubules and motors

Nuclear movements in muscle cells depend on the microtubule cytoskeleton and associated motor proteins (Cadot et al., 2012; Englander & Rubin, 1987; Wilson & Holzbaur, 2012). To determine whether the in vitro system could recapitulate this aspect of muscle cell organization, we imaged myotube nuclei in live squashes of interphase egg extract supplemented with rhodamine tubulin and the Hoechst DNA dye. We observed that nuclei moved apart at rates averaging 1-3 µm/min and that movement was completely lost upon addition of the microtubule-depolymerizing drug nocodazole (Figure 4A and B, and Supporting Information Videos 1 and 2). Treatment with the dynein inhibitor ciliobrevin also strongly impaired nuclear movement (Figure 4C and D, Supporting Information Videos 3 and 4) (Firestone et al., 2012). These results are consistent with the observed roles of microtubules and dynein in nuclear movement in muscle cells (Cadot et al., 2012; Wilson & Holzbaur, 2012). Interestingly, treatment with monastrol, an inhibitor of the kinesin 5 motor Eg5, increased the average rate of nuclear movement ∼2 fold, to 2.97±1.74 µm/min compared to 1.429 ±1.51 µm/min in control extracts (Figure 4C and D) (Mayer et al., 1999). However, the basis of this effect is unclear, since monastrol treatment resulted in the formation of more robust perinuclear microtubule networks (Supporting Information Video 5), and Eg5 has been shown to affect microtubule dynamics (Chen et al., 2017; Fridman et al., 2013; Gardner et al., 2008; Kapoor et al., 2000; Sharp et al., 1999).

**Figure 4.**
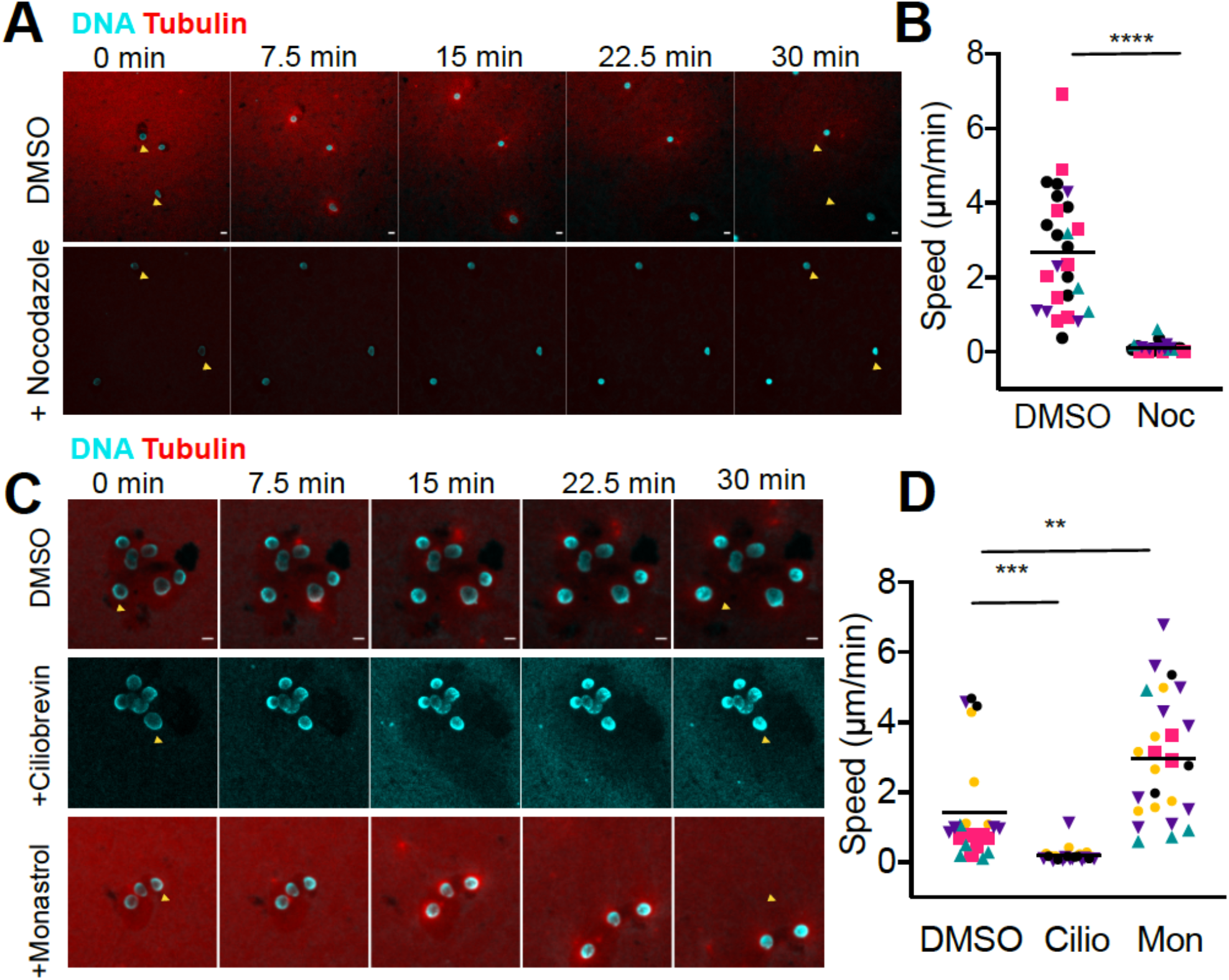
Nuclear movement in *Xenopus* egg extract is microtubule- and motor-dependent. (A) Image montage from 30 minute time-lapse videos of myotube nuclei in thin squashes of *Xenopus* egg extract showing nuclear movements are completely blocked in extract treated with 5 µg/mL of the microtubule depolymerizing drug nocodazole compared to the solvent control (DMSO). Yellow triangles indicate original positions of nuclei at t=0. (B) Quantification of rates showing that nuclear movement in control extract, 2.69±1.6 µm/min, is reduced to 0.11±1.21 µm/min upon nocodazole treatment, p <0.0001. Data pooled from n=4 extracts are depicted along with the mean. Individually color-coded points indicate rates from a single extract. (C) Image montage from 30 minute time lapse movies of myotube nuclei in extracts in the presence of solvent control, 25 µM dynein inhibitor ciliobrevin, or 25 µM Eg5 inhibitor monastrol. (D) Quantification of motility rates showing that dynein inhibition decreases nuclear movements from 1.43±1.51 µm/min to 0.193±0.21 µm/min (p = 0.0008) while Eg5 inhibition increases the rate of nuclear movement 2-fold (2.97± 1.74 µm/min, p = 0.0018). Scale bars are 10 µm.

Thus, live imaging of the in vitro assay allowed us to observe how myotube nuclei move and distribute themselves in a microtubule- and motor-dependent manner. However, the measured rates of movement were significantly higher than in cultured C2C12 myotubes (0.213 µm/min) (Wilson & Holzbaur, 2012). This discrepancy may be explained by the physically unconstrained nature of live squash preparations compared to muscle cells that are only tens of micrometers wide and contain ordered actomyosin arrays that could impede nuclear movements. Moreover, microtubules may be more dynamic in egg extract compared to the stable arrays found in muscle cells (Gundersen et al., 1989). The nuclear movements observed required adsorption of microtubules and motors to the coverslip. When coverslips were passivated, nuclei movement appeared to be driven by bulk extract flow (Supplemental Figure S2). Thus, nuclear movements may require buttressing of microtubule arrays against stable structures both in vitro and in vivo, providing a mechanism to evenly position nuclei at the periphery of the muscle fiber (Folker et al., 2012; William Roman & Gomes, 2018).

In conclusion, by combining features of mouse C2C12 cells with the biochemically tractable *Xenopus* egg extract we have developed an assay to identify and dissect the roles of different factors in perinuclear microtubule nucleation and subsequent nuclear movements. In vivo, however, microtubules emanate not only from the perinuclear region, but also from other cytoplasmic sites in muscle cells to coordinate several important functions, such as movement of myonuclei to sites of muscle injury (Bugnard et al., 2005; Gimpel et al., 2017; Musa et al., 2003; Oddoux et al., 2013; W Roman et al., 2021). It will be interesting to learn how different populations of microtubules in muscle cells contribute to this important process.

A major function of perinuclear microtubules could be to help organize subcellular domains in muscle cells, a long-standing idea in the field positing that each nucleus could be responsible for the fate of the cytoplasm in its immediate vicinity. In support of this hypothesis, subcellular localization of mRNA to the area surrounding source nuclei has been observed (Hall & Ralston, 1989; Pavlath et al., 1989; Ralston & Hall, 1989, 1992; Rotundo, 1990; Windner et al., 2019). This observation was recently reinforced, and for the first time, microtubules were implicated in a size-dependent sorting and distribution of mRNA molecules in muscle (Pinheiro et al., 2021). This would be consistent with the spatial restriction of proteins and mRNA that has been discovered in many other systems (Blower, 2013; Saxton, 2001).

Perinuclear ncMTOCs have also been identified in other cell types including *Drosophila* fat body cells and transiently during *Drosophila* oogenesis and it would be interesting to observe the subcellular organization that perinuclear microtubules mediate in these systems (Sun et al., 2019; Tillery et al., 2018; Zheng et al., 2020). An important open question is whether the same set of molecules organize the perinuclear ncMTOC in different of cell types or even among nuclei in different regions within a single muscle cell syncytium, for example near the neuromuscular junction. Further development of in vitro assays promises to shed light on how microtubules emanating or anchored at the perinuclear ncMTOC contribute to cellular organization in different tissue types.

## MATERIALS AND METHODS

### Animal models

*Xenopus laevis* frogs were maintained in accordance with the Animal Use and Care Protocol at University of California, Berkeley. Frogs were obtained from NASCO and were maintained in a recirculating system as previously described (Miller et al., 2019).

### *Xenopus* egg extract preparation

CSF-arrested egg extract was made as previously described (Hannak & Heald, 2006; Maresca & Heald, 2006). Briefly, eggs arrested in meiosis II of the cell cycle were collected, dejellied, packed and lysed by centrifugation. The cytoplasmic layer was collected using a syringe and supplemented with 10 mg/ml of leupeptin (Neta Scientific), pepstatin (Sigma Aldrich) and chymostatin (Millipore Sigma) (LPC), 20 mM of cytochalasin B (diluted 1:500) (cytoB - Sigma-Aldrich C6762) and energy mix (3.75 mM creatine phosphate, 0.5 mM ATP, 0.5 mM MgCl_2_, 0.05 mM EGTA). Extract was supplemented with 750 nM fluorescently labeled rhodamine tubulin. For live imaging, extract was also supplemented with 50 ng/ml Hoechst 33342 (Thermo Fisher) to label DNA.

### Cell culture and differentiation

Mouse myoblast C2C12 cells (ATCC Cat# CRL-1772, RRID:CVCL_0188) were obtained from the UC Berkeley cell culture facility and were cultured in DMEM (Gibco-Thermo Fisher 1165092 high glucose, +L-glutamine,10% FBS, with the addition of 100 U/ml Pen/Strep-Gibco). Cells were seeded at 5000 cells/ cm^2^ for passaging. Undifferentiated cultures were not allowed to exceed 60% confluence. Frozen cell stocks were thawed for each preparation of nuclei to ensure the myoblast population was not depleted. To induce differentiation, cells were allowed to reach confluence and media was switched with DMEM containing 2% horse serum (Invitrogen). Media was switched every day for 4 full days, and cells were harvested by trypsinization (0.25% Trypsin-EDTA, Gibco) for 1-2 min, to ensure only differentiated cells detached. Trypsinization was monitored by phase contrast microscopy. Purification of C2C12 cell nuclei was carried out as described with minor modifications (Srsen et al., 2009). Trypsin was neutralized and detached cells were washed 2x in PBS. The washed pellet was resuspended in a minimal volume of cold PBS to which was added an equal volume of hypotonic buffer (10 mM KCl, 10 mM PIPES pH 7.4, 0.5 mM MgCl_2_, 20 µM cytoB, 0.2 mM PMSF and 20 µg/ml LPC) and 0.75 volumes of PBS with 0.1% NP40. Subsequent steps were carried out quickly and on ice. The cell suspension was passed ∼3-4 times through a syringe and 27-gauge needle. Lysis was monitored by phase contrast microscopy. The material was layered over a 30% sucrose cushion in homogenization buffer (0.2 M KCl, 50 mM PIPES pH 7.4, 0.5 mM MgCl_2_, 0.1 mM PMSF, 10 µg/ml LPC). The nuclear pellet was stored in 50% glycerol, 250 mM sucrose, 80 mM KCl, 20 mM NaCl, 5 mM EGTA, 15 mM PIPES pH 7.4, 1 mM DTT, 0.5 mM spermidine, 0.2 mM spermine and protease inhibitors. For experiments comparing undifferentiated cell nuclei and nuclei isolated from cells on various days of differentiation, nuclei were isolated as described (Nabbi & Riabowol, 2015).

### Extract reactions

To induce interphase 1x Ca^+2^ solution was added from a 50X stock solution (20 mM CaCl_2_, 500 mM KCl, 5 mM MgCl_2_). C2C12 myotube (760 nuclei/µl of extract) or *Xenopus* sperm nuclei (1000 nuclei/µl - prepared as previously described) were added to extract 1 minute after the addition of 1x Ca^+2^ solution (Murray, 1991). Microtubules were allowed to assemble for 10 minutes after which 3.5 µl of the reaction was fixed by addition to an equal volume of fresh spindle fix (48% glycerol, 11% formaldehyde, 1 µg/ml Hoechst). Small molecules/ proteins were added to extracts as follows: 1 µM MLN8237 (Sigma-Aldrich) (or 0.5% DMSO) was added to CSF extract for a 10 minute preincubation. RNase-A (Sigma-Aldrich 10109142001) was added to a final concentration of 100 µg/ml for 1 hour prior to induction of interphase. Motor inhibitors (Ciliobrevin-Millipore 250401 and Monastrol -Sigma M8515) were added 10 minutes prior to interphase induction to a final concentration of 25 µM.

### Preparation of clean coverslips and live imaging of nuclei

Coverslips were sonicated for 30 minutes each in 1% Hellmanex detergent (Hellma USA Inc.) and 1 M KOH followed each by 10 washes in deionized water. Coverslips were nutated in 100% acetone, washed and sonicated in 100% ethanol and stored in ethanol as described with modifications (Kueh et al., 2008). Coverslips were dried with compressed air and a chamber was made by sandwiching 6.5 (for Fig. 4C,D) or 8 µl (for Fig 4A,B) of extract containing nuclei between two coverslips stuck to a glass slide with double-sided tape. Coverslips were passivated using PLL-PEG (SuSoS surface tech.) when applicable as described with modifications (dried with air instead of nitrogen) (Field et al., 2017). The live squash sandwich was sealed with VALAP. Time lapse images were obtained every 90s for 30 minutes.

### C2C12 nuclei immunofluorescence

Nuclei in egg extracts were mixed with 20 volumes fix buffer (ELB - 250 mM sucrose, 50 mM KCl, 2.5 mM MgCl_2_, and 10 mM HEPES pH 7.8, 15% glycerol, 3.7% formaldehyde) for 15 minutes at room temperature, layered over a 5 ml cushion buffer (1x XB buffer -10 mM HEPES pH 7.4, 100 mM KCl, 1 mM MgCl_2_, 0.1 mM CaCl_2_ with 200 mM sucrose & 25% glycerol), and spun onto 12 mm circular coverslips at 1000 x g for 15 minutes at 16°C. Nuclei were post-fixed in methanol (−20°C) for 1 minute and rehydrated in 1x PBS with 0.1% NP40. Coverslips were blocked with 3% Bovine Serum Albumin (BSA) in PBS for 1 hour at room temperature and incubated at room temperature for 1 hour each with primary and secondary antibody diluted in PBS-BSA followed by 1 µg/ml Hoechst, mounted in Vectashield (Vector Laboratories), and sealed with nail polish (Sally Hansen). Antibodies used were pericentrin (PCNT) antibody (PRB-432C-Biolegend, Covance Cat# PRB-432C, RRID:AB_2313709), GM130 (anti-rat amino acids 869-982 BD Transduction Laboratories), mouse monoclonal alpha-tubulin (Sigma-Aldrich Cat# T9026, RRID:AB_477593) and secondary antibodies goat anti mouse 488 and anti-rabbit 568 antibodies (Invitrogen).

### Microscopy and data quantification

Microscopes used were Olympus BX51 microscope with a Hamamatsu ORCA-ER camera. Objectives used were Olympus UPlan FL 20x/NA 0.5, 40x/NA 0.75 air, and Olympus PlanApo N 60x/NA 1.42 oil. Images and movies were taken with micromanager or Olympus CellSens Dimension 2 software (Edelstein et al., 2014). Confocal images were acquired on an inverted Zeiss LSM 800 confocal microscope with Zeiss PlanApo [20x/NA 0.8, 63x/NA 1.4 oil] objectives. In most cases, the average gray value corresponding to microtubule intensity in a doughnut shaped zone around the nucleus was measured (ie. total gray value/area of doughnut) from which the average gray value of background in each individual image was subtracted using a macro written in FIJI (Schindelin et al., 2012). For sperm asters, a circle of diameter 40 µm was used to encompass the center of the aster and average fluorescence was measured from which background was subtracted. At least 3 independent extracts were used for each experiment and a representative graph was depicted due to the nature of variability between *Xenopus* egg extracts. Data across extracts was pooled in Figure 4B & 4D (where conclusions were confirmed also by analyzing mean of means). Data was determined to follow a normal distribution by a QQ plot. Control versus drug experiments were analyzed using t-test with Welch’s correction (for unequal variances) using GraphPad Prism version 8.2.1 for Mac, GraphPad Software, San Diego, California USA, www.graphpad.com. For nuclear movement, with control and drug-treated extracts, distance traveled was obtained by using the Trackmate plugin divided by total time of the track (Tinevez et al., 2017). Linear scaling of microtubule fluorescence between compared images was always the same if fluorescence was to be quantified. In exceptional cases where Hoechst (DNA) signal varied greatly due to squash conditions, fluorescence intensity was scaled to match visually as quantitative comparisons of DNA were not carried out in this work (Figure 4C).

## ACKNOWLEDGMENTS

We thank Maiko Kitaoka for help with microscopy, Christian Erikson for help with data storage, and rotation students Claire Goul and Cynthia Harris for their assistance with experiments. Thank you to the Canzio and Buchwalter labs at UCSF and the Hexalab at UCB for discussion, and thanks to Maiko Kitaoka, Clotilde Cadart and Helena Cantwell for critical reading of the manuscript. AVN was supported by the American Heart Association AWRP fellowship 17POST33670285. RH was supported by NIH R35GM118183 and the Flora Lamson Hewlett chair.

## AUTHOR CONTRIBUTIONS

A.V.N. performed the experiments, analyzed the data and prepared the figures. A.V.N. and R.H. wrote the manuscript.

## REFERENCES

Abmayr, S. M., & Pavlath, G. K. (2012). Myoblast fusion: lessons from flies and mice. Development, 139(4), 641–656. https://doi.org/10.1242/dev.068353

Apel, E. D., Lewis, R. M., Grady, R. M., & Sanes, J. R. (2000). Syne-1, a dystrophin-and Klarsicht-related protein associated with synaptic nuclei at the neuromuscular junction. The Journal of Biological Chemistry, 275(41), 31986–31995. https://doi.org/10.1074/jbc.M004775200

Azevedo, M., & Baylies, M. K. (2020). Getting into Position: Nuclear Movement in Muscle Cells. Trends in Cell Biology, 30(4), 303–316. https://doi.org/10.1016/j.tcb.2020.01.002

Balczon, R., Bao, L., & Zimmer, W. E. (1994). PCM-1, A 228-kD centrosome autoantigen with a distinct cell cycle distribution. The Journal of Cell Biology, 124(5), 783–793. https://doi.org/10.1083/jcb.124.5.783

Becker, R., Leone, M., & Engel, F. B. (2020). Microtubule organization in striated muscle cells. Cells, 9(6). https://doi.org/10.3390/cells9061395

Blower, M. D. (2013). Molecular insights into intracellular RNA localization. International Review of Cell and Molecular Biology, 302, 1–39. https://doi.org/10.1016/B978-0-12-407699-0.00001-7

Blower, M. D., Nachury, M., Heald, R., & Weis, K. (2005). A Rae1-containing ribonucleoprotein complex is required for mitotic spindle assembly. Cell, 121(2), 223–234. https://doi.org/10.1016/j.cell.2005.02.016

Bugnard, E., Zaal, K. J. M., & Ralston, E. (2005). Reorganization of microtubule nucleation during muscle differentiation. Cell Motility and the Cytoskeleton, 60(1), 1–13. https://doi.org/10.1002/cm.20042

Cadot, B., Gache, V., Vasyutina, E., Falcone, S., Birchmeier, C., & Gomes, E. R. (2012). Nuclear movement during myotube formation is microtubule and dynein dependent and is regulated by Cdc42, Par6 and Par3. EMBO Reports, 13(8), 741–749. https://doi.org/10.1038/embor.2012.89

Caporizzo, M. A., Chen, C. Y., Salomon, A. K., Margulies, K. B., & Prosser, B. L. (2018). Microtubules provide a viscoelastic resistance to myocyte motion. Biophysical Journal, 115(9), 1796–1807. https://doi.org/10.1016/j.bpj.2018.09.019

Cavazza, T., & Vernos, I. (2015). The RanGTP Pathway: From Nucleo-Cytoplasmic Transport to Spindle Assembly and Beyond. Frontiers in Cell and Developmental Biology, 3, 82. https://doi.org/10.3389/fcell.2015.00082

Chabin-Brion, K., Marceiller, J., Perez, F., Settegrana, C., Drechou, A., Durand, G., & Poüs, C. (2001). The Golgi complex is a microtubule-organizing organelle. Molecular Biology of the Cell, 12(7), 2047–2060. https://doi.org/10.1091/mbc.12.7.2047

Chen, G.-Y., Kang, Y. J., Gayek, A. S., Youyen, W., Tüzel, E., Ohi, R., & Hancock, W. O. (2017). Eg5 inhibitors have contrasting effects on microtubule stability and metaphase spindle integrity. ACS Chemical Biology, 12(4), 1038–1046. https://doi.org/10.1021/acschembio.6b01040

Cheng, X., & Ferrell, J. E. (2019). Spontaneous emergence of cell-like organization in Xenopus egg extracts. Science, 366(6465), 631–637. https://doi.org/10.1126/science.aav7793

Dammermann, A., & Merdes, A. (2002). Assembly of centrosomal proteins and microtubule organization depends on PCM-1. The Journal of Cell Biology, 159(2), 255–266. https://doi.org/10.1083/jcb.200204023

Dictenberg, J. B., Zimmerman, W., Sparks, C. A., Young, A., Vidair, C., Zheng, Y., Carrington, W., Fay, F. S., & Doxsey, S. J. (1998). Pericentrin and gamma-tubulin form a protein complex and are organized into a novel lattice at the centrosome. The Journal of Cell Biology, 141(1), 163–174. https://doi.org/10.1083/jcb.141.1.163

Doxsey, S. J., Stein, P., Evans, L., Calarco, P. D., & Kirschner, M. (1994). Pericentrin, a highly conserved centrosome protein involved in microtubule organization. Cell, 76(4), 639–650. https://doi.org/10.1016/0092-8674(94)90504-5

Duong, N. T., Morris, G. E., Lam, L. T., Zhang, Q., Sewry, C. A., Shanahan, C. M., & Holt, I. (2014). Nesprins: tissue-specific expression of epsilon and other short isoforms. Plos One, 9(4), e94380. https://doi.org/10.1371/journal.pone.0094380

Edelstein, A. D., Tsuchida, M. A., Amodaj, N., Pinkard, H., Vale, R. D., & Stuurman, N. (2014). Advanced methods of microscope control using μManager software. Journal of Biological Methods, 1(2). https://doi.org/10.14440/jbm.2014.36

Englander, L. L., & Rubin, L. L. (1987). Acetylcholine receptor clustering and nuclear movement in muscle fibers in culture. The Journal of Cell Biology, 104(1), 87–95. https://doi.org/10.1083/jcb.104.1.87

Espigat-Georger, A., Dyachuk, V., Chemin, C., Emorine, L., & Merdes, A. (2016). Nuclear alignment in myotubes requires centrosome proteins recruited by nesprin-1. Journal of Cell Science, 129(22), 4227–4237. https://doi.org/10.1242/jcs.191767

Fant, X., Srsen, V., Espigat-Georger, A., & Merdes, A. (2009). Nuclei of non-muscle cells bind centrosome proteins upon fusion with differentiating myoblasts. Plos One, 4(12), e8303. https://doi.org/10.1371/journal.pone.0008303

Field, C. M., Pelletier, J. F., & Mitchison, T. J. (2017). Xenopus extract approaches to studying microtubule organization and signaling in cytokinesis. Methods in Cell Biology, 137, 395– 435. https://doi.org/10.1016/bs.mcb.2016.04.014

Firestone, A. J., Weinger, J. S., Maldonado, M., Barlan, K., Langston, L. D., O’Donnell, M., Gelfand, V. I., Kapoor, T. M., & Chen, J. K. (2012). Small-molecule inhibitors of the AAA+ ATPase motor cytoplasmic dynein. Nature, 484(7392), 125–129. https://doi.org/10.1038/nature10936

Folker, E. S., Schulman, V. K., & Baylies, M. K. (2012). Muscle length and myonuclear position are independently regulated by distinct Dynein pathways. Development, 139(20), 3827–3837. https://doi.org/10.1242/dev.079178

Fridman, V., Gerson-Gurwitz, A., Shapira, O., Movshovich, N., Lakämper, S., Schmidt, C. F., & Gheber, L. (2013). Kinesin-5 Kip1 is a bi-directional motor that stabilizes microtubules and tracks their plus-ends in vivo. Journal of Cell Science, 126(Pt 18), 4147–4159. https://doi.org/10.1242/jcs.125153

Gardner, M. K., Bouck, D. C., Paliulis, L. V., Meehl, J. B., O’Toole, E. T., Haase, J., Soubry, A., Joglekar, A. P., Winey, M., Salmon, E. D., Bloom, K., & Odde, D. J. (2008). Chromosome congression by Kinesin-5 motor-mediated disassembly of longer kinetochore microtubules. Cell, 135(5), 894–906. https://doi.org/10.1016/j.cell.2008.09.046

Gimpel, P., Lee, Y. L., Sobota, R. M., Calvi, A., Koullourou, V., Patel, R., Mamchaoui, K., Nédélec, F., Shackleton, S., Schmoranzer, J., Burke, B., Cadot, B., & Gomes, E. R. (2017). Nesprin-1α-Dependent Microtubule Nucleation from the Nuclear Envelope via Akap450 Is Necessary for Nuclear Positioning in Muscle Cells. Current Biology, 27(19), 2999–3009.e9. https://doi.org/10.1016/j.cub.2017.08.031

Gundersen, G. G., Khawaja, S., & Bulinski, J. C. (1989). Generation of a stable, posttranslationally modified microtubule array is an early event in myogenic differentiation. The Journal of Cell Biology, 109(5), 2275–2288. https://doi.org/10.1083/jcb.109.5.2275

Hall, Z. W., & Ralston, E. (1989). Nuclear domains in muscle cells. Cell, 59(5), 771–772. https://doi.org/10.1016/0092-8674(89)90597-7

Hannak, E., & Heald, R. (2006). Investigating mitotic spindle assembly and function in vitro using Xenopus laevis egg extracts. Nature Protocols, 1(5), 2305–2314. https://doi.org/10.1038/nprot.2006.396

Heffler, J., Shah, P. P., Robison, P., Phyo, S., Veliz, K., Uchida, K., Bogush, A., Rhoades, J., Jain, R., & Prosser, B. L. (2020). A balance between intermediate filaments and microtubules maintains nuclear architecture in the cardiomyocyte. Circulation Research, 126(3), e10–e26. https://doi.org/10.1161/CIRCRESAHA.119.315582

Henderson, C. A., Gomez, C. G., Novak, S. M., Mi-Mi, L., & Gregorio, C. C. (2017). Overview of the muscle cytoskeleton. Comprehensive Physiology, 7(3), 891–944. https://doi.org/10.1002/cphy.c160033

Ide, K., Muko, M., & Hayashi, K. (2021). The Golgi apparatus is the main microtubule-organizing center in differentiating skeletal muscle cells. Histochemistry and Cell Biology, 156(3), 273–281. https://doi.org/10.1007/s00418-021-01999-6

Ishihara, K., Nguyen, P. A., Groen, A. C., Field, C. M., & Mitchison, T. J. (2014). Microtubule nucleation remote from centrosomes may explain how asters span large cells. Proceedings of the National Academy of Sciences of the United States of America, 111(50), 17715–17722. https://doi.org/10.1073/pnas.1418796111

Kapoor, T. M., Mayer, T. U., Coughlin, M. L., & Mitchison, T. J. (2000). Probing spindle assembly mechanisms with monastrol, a small molecule inhibitor of the mitotic kinesin, Eg5. The Journal of Cell Biology, 150(5), 975–988. https://doi.org/10.1083/jcb.150.5.975

Kronebusch, P. J., & Singer, S. J. (1987). The microtubule-organizing complex and the Golgi apparatus are co-localized around the entire nuclear envelope of interphase cardiac myocytes. Journal of Cell Science, 88 (Pt 1), 25–34.

Kubo, A., Sasaki, H., Yuba-Kubo, A., Tsukita, S., & Shiina, N. (1999). Centriolar satellites: molecular characterization, ATP-dependent movement toward centrioles and possible involvement in ciliogenesis. The Journal of Cell Biology, 147(5), 969–980. https://doi.org/10.1083/jcb.147.5.969

Kueh, H. Y., Charras, G. T., Mitchison, T. J., & Brieher, W. M. (2008). Actin disassembly by cofilin, coronin, and Aip1 occurs in bursts and is inhibited by barbed-end cappers. The Journal of Cell Biology, 182(2), 341–353. https://doi.org/10.1083/jcb.200801027

Lu, Z., Joseph, D., Bugnard, E., Zaal, K. J., & Ralston, E. (2001). Golgi complex reorganization during muscle differentiation: visualization in living cells and mechanism. Molecular Biology of the Cell, 12(4), 795–808. https://doi.org/10.1091/mbc.12.4.795

Magnaghi-Jaulin, L., Eot-Houllier, G., Gallaud, E., & Giet, R. (2019). Aurora A protein kinase: to the centrosome and beyond. Biomolecules, 9(1). https://doi.org/10.3390/biom9010028

Maresca, T. J., & Heald, R. (2006). Methods for studying spindle assembly and chromosome condensation in Xenopus egg extracts. Methods in Molecular Biology, 322, 459–474. https://doi.org/10.1007/978-1-59745-000-3_33

Mayer, T. U., Kapoor, T. M., Haggarty, S. J., King, R. W., Schreiber, S. L., & Mitchison, T. J. (1999). Small molecule inhibitor of mitotic spindle bipolarity identified in a phenotype-based screen. Science, 286(5441), 971–974. https://doi.org/10.1126/science.286.5441.971

Miller, K. E., Session, A. M., & Heald, R. (2019). Kif2a Scales Meiotic Spindle Size in Hymenochirus boettgeri. Current Biology, 29(21), 3720–3727.e5. https://doi.org/10.1016/j.cub.2019.08.073

Murray, A. W. (1991). Cell cycle extracts. Methods in Cell Biology, 36, 581–605.

Musa, H., Orton, C., Morrison, E. E., & Peckham, M. (2003). Microtubule assembly in cultured myoblasts and myotubes following nocodazole induced microtubule depolymerisation. Journal of Muscle Research and Cell Motility, 24(4–6), 301–308.

Nabbi, A., & Riabowol, K. (2015). Rapid Isolation of Nuclei from Cells In Vitro. Cold Spring Harbor Protocols, 2015(8), 769–772. https://doi.org/10.1101/pdb.prot083733

Nguyen, P. A., Groen, A. C., Loose, M., Ishihara, K., Wühr, M., Field, C. M., & Mitchison, T. J. (2014). Spatial organization of cytokinesis signaling reconstituted in a cell-free system. Science, 346(6206), 244–247. https://doi.org/10.1126/science.1256773

Oddoux, S., Zaal, K. J., Tate, V., Kenea, A., Nandkeolyar, S. A., Reid, E., Liu, W., & Ralston, E. (2013). Microtubules that form the stationary lattice of muscle fibers are dynamic and nucleated at Golgi elements. The Journal of Cell Biology, 203(2), 205–213. https://doi.org/10.1083/jcb.201304063

Pavlath, G. K., Rich, K., Webster, S. G., & Blau, H. M. (1989). Localization of muscle gene products in nuclear domains. Nature, 337(6207), 570–573. https://doi.org/10.1038/337570a0

Pinheiro, H., Pimentel, M. R., Sequeira, C., Oliveira, L. M., Pezzarossa, A., Roman, W., & Gomes, E. R. (2021). mRNA distribution in skeletal muscle is associated with mRNA size. Journal of Cell Science, 134(14). https://doi.org/10.1242/jcs.256388

Ralston, E. (1993). Changes in architecture of the Golgi complex and other subcellular organelles during myogenesis. The Journal of Cell Biology, 120(2), 399–409. https://doi.org/10.1083/jcb.120.2.399

Ralston, E., & Hall, Z. W. (1989). Transfer of a protein encoded by a single nucleus to nearby nuclei in multinucleated myotubes. Science, 244(4908), 1066–1069. https://doi.org/10.1126/science.2543074

Ralston, E., & Hall, Z. W. (1992). Restricted distribution of mRNA produced from a single nucleus in hybrid myotubes. The Journal of Cell Biology, 119(5), 1063–1068. https://doi.org/10.1083/jcb.119.5.1063

Randles, K. N., Lam, L. T., Sewry, C. A., Puckelwartz, M., Furling, D., Wehnert, M., McNally, E. M., & Morris, G. E. (2010). Nesprins, but not sun proteins, switch isoforms at the nuclear envelope during muscle development. Developmental Dynamics, 239(3), 998–1009. https://doi.org/10.1002/dvdy.22229

Robison, P., Caporizzo, M. A., Ahmadzadeh, H., Bogush, A. I., Chen, C. Y., Margulies, K. B., Shenoy, V. B., & Prosser, B. L. (2016). Detyrosinated microtubules buckle and bear load in contracting cardiomyocytes. Science, 352(6284), aaf0659. https://doi.org/10.1126/science.aaf0659

Roman, W, Pinheiro, H., Pimentel, M. R., Segalés, J., Oliveira, L. M., García-Domínguez, E., Gómez-Cabrera, M. C., Serrano, A. L., Gomes, E. R., & Muñoz-Cánoves, P. (2021). Muscle repair after physiological damage relies on nuclear migration for cellular reconstruction. Science, 374(6565), 355–359. https://doi.org/10.1126/science.abe5620

Roman, William, & Gomes, E. R. (2018). Nuclear positioning in skeletal muscle. Seminars in Cell & Developmental Biology, 82, 51–56. https://doi.org/10.1016/j.semcdb.2017.11.005

Rotundo, R. L. (1990). Nucleus-specific translation and assembly of acetylcholinesterase in multinucleated muscle cells. The Journal of Cell Biology, 110(3), 715–719. https://doi.org/10.1083/jcb.110.3.715

Sanchez, A. D., & Feldman, J. L. (2017). Microtubule-organizing centers: from the centrosome to non-centrosomal sites. Current Opinion in Cell Biology, 44, 93–101. https://doi.org/10.1016/j.ceb.2016.09.003

Saxton, W. M. (2001). Microtubules, Motors, and mRNA Localization Mechanisms. Cell, 107(6), 707–710. https://doi.org/10.1016/S0092-8674(01)00602-X

Schindelin, J., Arganda-Carreras, I., Frise, E., Kaynig, V., Longair, M., Pietzsch, T., Preibisch, S., Rueden, C., Saalfeld, S., Schmid, B., Tinevez, J.-Y., White, D. J., Hartenstein, V., Eliceiri, K., Tomancak, P., & Cardona, A. (2012). Fiji: an open-source platform for biological-image analysis. Nature Methods, 9(7), 676–682. https://doi.org/10.1038/nmeth.2019

Schmidt, P. H., Dransfield, D. T., Claudio, J. O., Hawley, R. G., Trotter, K. W., Milgram, S. L., & Goldenring, J. R. (1999). AKAP350, a multiply spliced protein kinase A-anchoring protein associated with centrosomes. The Journal of Biological Chemistry, 274(5), 3055–3066. https://doi.org/10.1074/jbc.274.5.3055

Sepulveda, G., Antkowiak, M., Brust-Mascher, I., Mahe, K., Ou, T., Castro, N. M., Christensen, L. N., Cheung, L., Jiang, X., Yoon, D., Huang, B., & Jao, L.-E. (2018). Co-translational protein targeting facilitates centrosomal recruitment of PCNT during centrosome maturation in vertebrates. ELife, 7. https://doi.org/10.7554/eLife.34959

Sharp, D. J., McDonald, K. L., Brown, H. M., Matthies, H. J., Walczak, C., Vale, R. D., Mitchison, T. J., & Scholey, J. M. (1999). The bipolar kinesin, KLP61F, cross-links microtubules within interpolar microtubule bundles of Drosophila embryonic mitotic spindles. The Journal of Cell Biology, 144(1), 125–138. https://doi.org/10.1083/jcb.144.1.125

Srsen, V., Fant, X., Heald, R., Rabouille, C., & Merdes, A. (2009). Centrosome proteins form an insoluble perinuclear matrix during muscle cell differentiation. BMC Cell Biology, 10, 28. https://doi.org/10.1186/1471-2121-10-28

Sun, T., Song, Y., Dai, J., Mao, D., Ma, M., Ni, J.-Q., Liang, X., & Pastor-Pareja, J. C. (2019). Spectraplakin shot maintains perinuclear microtubule organization in drosophila polyploid cells. Developmental Cell, 49(5), 731–747.e7. https://doi.org/10.1016/j.devcel.2019.03.027

Takahashi, M., Shibata, H., Shimakawa, M., Miyamoto, M., Mukai, H., & Ono, Y. (1999). Characterization of a novel giant scaffolding protein, CG-NAP, that anchors multiple signaling enzymes to centrosome and the golgi apparatus. The Journal of Biological Chemistry, 274(24), 17267–17274. https://doi.org/10.1074/jbc.274.24.17267

Tassin, A. M., Maro, B., & Bornens, M. (1985). Fate of microtubule-organizing centers during myogenesis in vitro. The Journal of Cell Biology, 100(1), 35–46. https://doi.org/10.1083/jcb.100.1.35

Tillery, M. M. L., Blake-Hedges, C., Zheng, Y., Buchwalter, R. A., & Megraw, T. L. (2018). Centrosomal and Non-Centrosomal Microtubule-Organizing Centers (MTOCs) in Drosophila melanogaster. Cells, 7(9). https://doi.org/10.3390/cells7090121

Tinevez, J.-Y., Perry, N., Schindelin, J., Hoopes, G. M., Reynolds, G. D., Laplantine, E., Bednarek, S. Y., Shorte, S. L., & Eliceiri, K. W. (2017). TrackMate: An open and extensible platform for single-particle tracking. Methods, 115, 80–90. https://doi.org/10.1016/j.ymeth.2016.09.016

Tsai, M.-Y., & Zheng, Y. (2005). Aurora A kinase-coated beads function as microtubule-organizing centers and enhance RanGTP-induced spindle assembly. Current Biology, 15(23), 2156–2163. https://doi.org/10.1016/j.cub.2005.10.054

Warren, R. H. (1974). Microtubular organization in elongating myogenic cells. The Journal of Cell Biology, 63(2 Pt 1), 550–566. https://doi.org/10.1083/jcb.63.2.550

Wilson, M. H., & Holzbaur, E. L. F. (2012). Opposing microtubule motors drive robust nuclear dynamics in developing muscle cells. Journal of Cell Science, 125(Pt 17), 4158–4169. https://doi.org/10.1242/jcs.108688

Wilson, M. H., & Holzbaur, E. L. F. (2015). Nesprins anchor kinesin-1 motors to the nucleus to drive nuclear distribution in muscle cells. Development, 142(1), 218–228. https://doi.org/10.1242/dev.114769

Windner, S. E., Manhart, A., Brown, A., Mogilner, A., & Baylies, M. K. (2019). Nuclear Scaling Is Coordinated among Individual Nuclei in Multinucleated Muscle Fibers. Developmental Cell, 49(1), 48–62.e3. https://doi.org/10.1016/j.devcel.2019.02.020

Witczak, O., Skålhegg, B. S., Keryer, G., Bornens, M., Taskén, K., Jahnsen, T., & Orstavik, S. (1999). Cloning and characterization of a cDNA encoding an A-kinase anchoring protein located in the centrosome, AKAP450. The EMBO Journal, 18(7), 1858–1868. https://doi.org/10.1093/emboj/18.7.1858

Zhang, Q., Skepper, J. N., Yang, F., Davies, J. D., Hegyi, L., Roberts, R. G., Weissberg, P. L., Ellis, J. A., & Shanahan, C. M. (2001). Nesprins: a novel family of spectrin-repeat-containing proteins that localize to the nuclear membrane in multiple tissues. Journal of Cell Science, 114(Pt 24), 4485–4498. https://doi.org/10.1242/jcs.114.24.4485

Zheng, Y., Buchwalter, R. A., Zheng, C., Wight, E. M., Chen, J. V., & Megraw, T. L. (2020). A perinuclear microtubule-organizing centre controls nuclear positioning and basement membrane secretion. Nature Cell Biology, 22(3), 297–309. https://doi.org/10.1038/s41556-020-0470-7

