## Supplementary figures and video legends for "Reconstitution of muscle cell microtubule organization in vitro"

**SUPPLEMENTARY DATA: 2 FIGURES AND 5 VIDEOS**

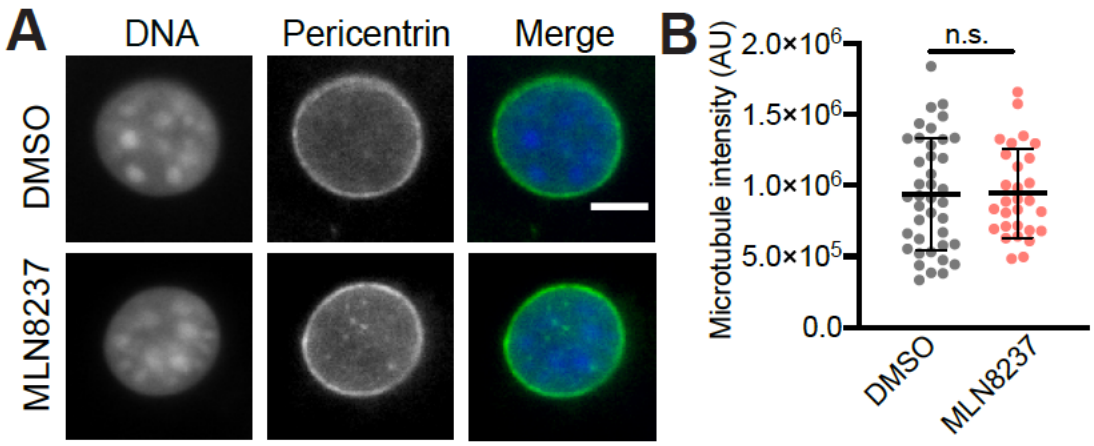


**Supplementary Figure S1. Aurora A inhibition does not affect pericentrin distribution around C2C12 nuclei.** (A) Representative images showing DNA (blue) and pericentrin (green) staining of nuclei treated with DMSO or 1 µM Aurora A inhibitor MLN8237 (B) Quantification of pericentrin fluorescence (representative graph from n=3 extracts) shows that pericentrin intensity is unaffected following inhibitor treatment. Scale bar is 5 µm.

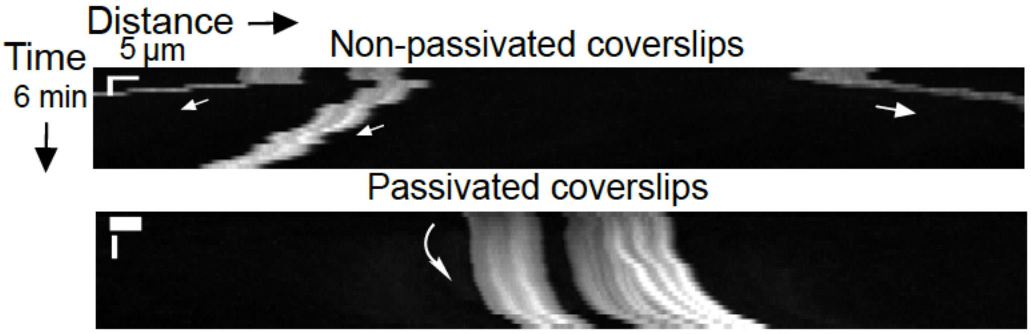


**Supplementary Figure S2**. **Unpassivated coverslips aid movement of nuclei in directions different from those of extract flow.** Kymographs of nuclei (x-axis: distance, bar = 5 µm, y-axis: time = 6 min) with arrowheads indicating the direction of movement of individual nuclei that appear as squiggly lines. Passivation causes all the nuclei to move in the same direction. Representative kymograph from n=3 extracts and 5 time-lapse movies.

**Supplementary Video S1.** Time lapse video of myotube nuclei in a live squash of interphase egg extract treated with 0.5% DMSO (solvent control). Images were captured every 90 seconds over a period of 30 minutes.

**Supplementary Video S2.** Time lapse video of myotube nuclei in a live squash of interphase egg extract treated with 5 μg/ml nocodazole to depolymerize microtubules.

**Supplementary Video S3.** Time lapse video of a cluster of myotube nuclei in egg extract treated with 0.5% DMSO alone over the course of 30 minutes.

**Supplementary Video S4.** Time lapse video of myotube nuclei in egg extract treated with 25 μM dynein inhibitor ciliobrevin.

**Supplementary Video S5.** Time lapse video of myotube nuclei in egg extract treated with 25 μM Eg5 inhibitor monastrol.
